# Shedding light on fenestrations in carnivorous pitcher plants: a test for convergent function

**DOI:** 10.64898/2026.09.20.751569

**Authors:** Emma S. Wagner, Sylvie Martin-Eberhardt, Marjorie Weber, Kadeem Gilbert

## Abstract

A longstanding question in evolution is if similar morphologies shared by distantly related organisms facing similar environmental pressures belie similar functions. The carnivorous pitcher form has evolved in three distinct plant lineages and encompasses a suite of convergent morphological traits, providing a natural experiment to test hypothesized links between morphology and function. Multiple pitcher plant species possess fenestrations (i.e., “small windows”) that let light pass through the pitcher leaf. While research on one species, *Nepenthes aristolochioides,* demonstrates that fenestrations manipulate light to create false exits and increase prey capture, whether this convergent morphology serves this convergent function in other pitcher species remains unresolved. We explored the function of fenestrations in two additional species of pitcher plants, *Nepenthes klossii* and *Cephalotus follicularis,* where the morphology and function of fenestration were previously unexplored. We inspected micromorphology of pitcher structure, conducted an insect bioassay with live plants to test for links between morphology and function, and quantified traits associated with light manipulation from herbarium specimens. Both species exhibited fenestration and related pitcher morphology similar to *N. aristolochioides.* Yet our experiments did not reveal a significant effect of light Author Contributions: ESW and SME conceived and designed the experiments, performed experiments, collected data, analyzed the data, and wrote the manuscript; other authors provided editorial advice. passing through fenestrations on fly capture for these two species. Key differences between these species and the previously investigated species may be the presence of wax crystals and the absence of viscoelastic fluid. While future studies may uncover weak to moderate light manipulation function not detected in our study, our data do not support convergent function in these distantly related, morphologically similar species.

## Introduction

Distantly related organisms exhibiting similar forms is a phenomenon that has captivated biologists since Darwin and raises several questions: Does similar morphology reveal analogous function? Are similar morphologies shaped by the same selective pressures acting on different lineages? When a convergent function necessitates a suite of component traits, how do those component traits evolve independently from one another? By comparing organisms with convergent morphologies, we can test for a convergent function and identify the requisite component traits.

One example of convergent evolution is carnivorous pitcher plants, which are an excellent system for investigating form following function and trait variances that allow complex function to arise. Carnivory acts as a nutrient supplement for plants in nutrient poor environments and involves attracting, trapping, and digesting prey through unique cup-shaped leaves (‘pitchers’). This specific prey capture mechanism has evolved independently in at least three orders of plants (Mithöfer 2011), but within this general morphology there are recurring traits whose function and benefit to the plant remains underexplored. One example of a convergent but relatively little studied phenotype of pitchers are fenestrations: thin, apigmented sections of the leaf where light can enter pitchers. The hypothesized function of fenestrations is to confuse potential prey inside the pitcher with a false exit and entrap them via manipulation of light; if light is concentrated coming through fenestrations opposite from the actual pitcher opening, arthropods may be misdirected. Previous work provided evidence that light passing through fenestrations significantly increased the amount of prey capture in *Nepenthes aristolochioides,* possibly by creating a false exit and confusing arthropods (Moran et al. 2012). The convergently evolved species *Sarracenia minor* has also been tested for light manipulation, although results in this species conflict (McGregor et al. 2016; Schaefer and Ruxton 2014), leaving the function of fenestrations in this and other untested pitcher plant species unresolved. The repeated evolution of fenestrations across convergently evolved carnivorous pitcher plants thus offers an opportunity to test for a convergent use of light manipulation linked to this specialized morphology.

In addition to fenestrations, we hypothesize several other component traits are required for light manipulation, each within a most effective morphological range. A small lid angle and small opening diameter could limit the amount of exterior light entering the pitcher, reduce the size of the true exit, and draw prey towards the light passing through the fenestrations. In addition, a concentration of fenestrations on the upper back wall of the pitcher opposite from the pitcher mouth would be required to place the false exit in one area where sunlight will pass through for much of the day. If a small lid angle, small opening diameter, and concentration of fenestration are acted upon by natural selection favoring light manipulation as a prey capture strategy, we would expect these traits to be in similar ranges to those of *N. aristolochioides*.

In this work, we explored the light manipulation function, what traits may work together for the function to occur, and whether those traits have a morphological range consistent with a light trapping function in two convergently evolved pitcher plants hypothesized to manipulate light for prey capture: *Nepenthes klossii* and *Cephalotus follicularis* (Moran et al. 2012; Thorogood et al. 2018). Despite being distantly related (Murphy et al. 2020), these species have strikingly similar morphology to both *N. aristolochioides* and *S. minor* (Figure 1). We used an insect bioassay with *C. follicularis* and *N. klossii* to test how prey capture is affected by light entering fenestrations. We complemented these bioassays with a phylogenetically paired quantification of component trait values of lid angle and opening diameter of herbarium specimens of species with and without light manipulating morphology. We asked: 1) Does the morphology of fenestration and proposed component traits (lid angle and opening diameter) quantitatively match phenotypes of species previously shown to use light manipulation in prey capture, consistent with multi-trait convergence? 2) Do associated pitcher traits display ranges consistent with a light manipulation function, as compared to closely related species without light manipulation? And finally, 3) does light manipulation through fenestrations impact prey capture for *N. klossii* and *C. follicularis*, consistent with convergent morphology and specialized function found in other pitcher plant species?

**Figure 1.**
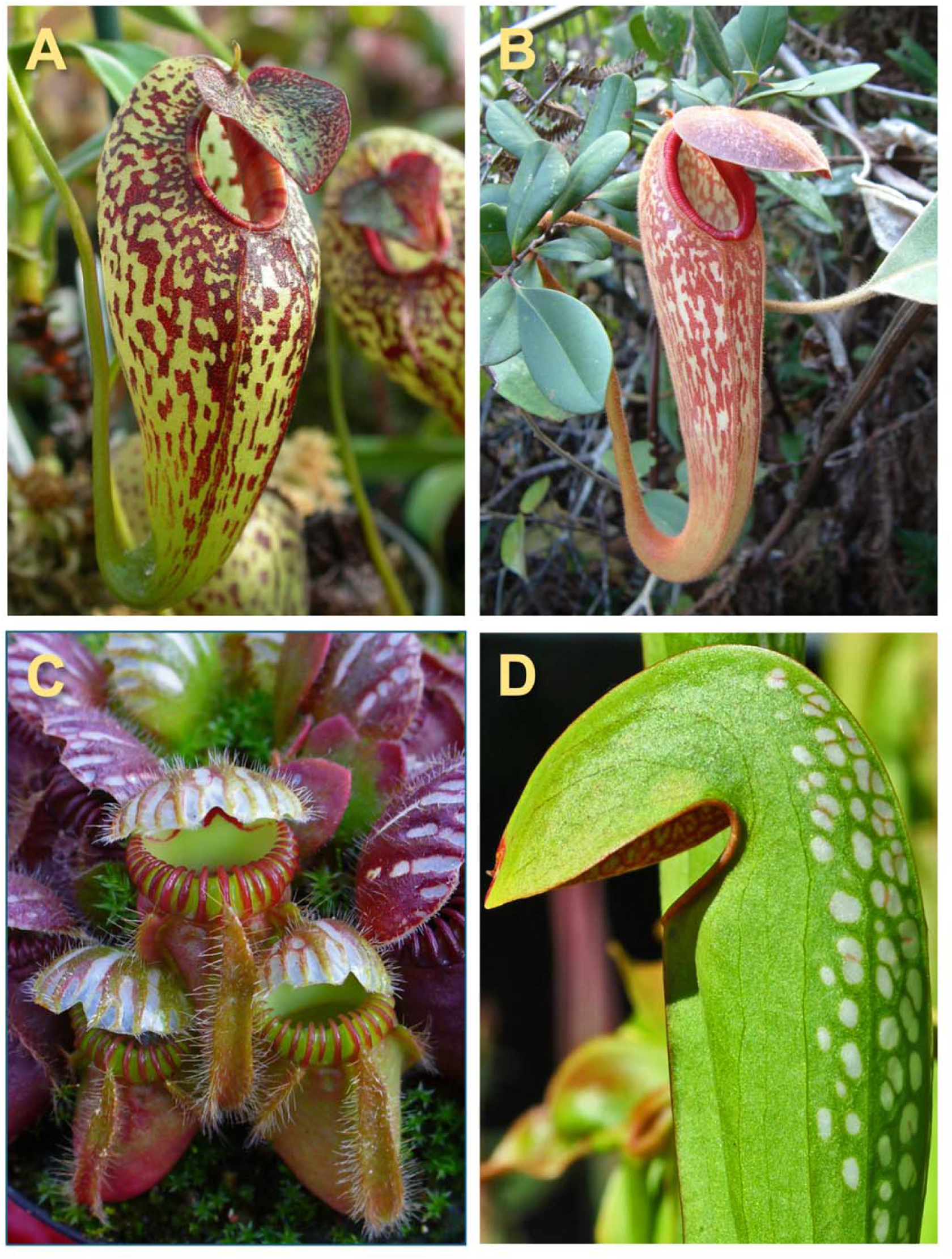
Fenestrations (apigmented leaf areas that let light through) on four convergently evolved pitcher plant species: A) *Nepenthes aristolochioides* B) *Nepenthes klossii* C) *Cephalotus follicularis* and D) *Sarrencia minor.* Photo credits: A) JeremiahsCPs, B) Alfindra Primaldh, CC BY 2.0, C) H. Zell CC BY-SA 3.0, D) Aaron Carlson CC BY-SA 2.0.

## Methods

### System

We focus on two phylogenetically and geographically distinct species of pitcher plants hypothesized to use light manipulation in prey capture: *N. klossii* and *C. follicularis. Nepenthes klossii* is a vining Asian pitcher plant found in Papua New Guinea. Despite being geographically and phylogenetically distant from *N. aristolochioides* within the *Nepenthes* genus*, N. klossii* has a strikingly similar mottled fenestration on its pitchers (Figure 1) and has been hypothesized to manipulate light to increase prey capture like *N. aristolochioides* (McPherson 2009; Moran et al. 2012). *Cephalotus follicularis,* also known as the Albany pitcher plant, is a small, low growing plant found in Australia in the monotypic family Cephalotaceae. This species has striped fenestrations on its lid that are distinctive from surrounding epidermis because of their whiteness and thinness (Figure 1). Fenestrations occur similarly in the Sarracenia family in combination with domed morphology, as seen in *S. minor* (Figure 1)*, S. psittacina,* and *Darlingtonia californica* (Thorogood et al. 2018). In this work we define fenestration as thin white areas of the pitcher as in seen in the Sarraceniaceae and Cephalotaceae families, as well as the lighter stripes that occur on mottled Nepenthaceae pitchers.

### Plant rearing

We grew four identical clones of both *N. klossii* (Carnivero, Austin, Texas) and *C. follicularis* (Curious Plant in Loveland, Ohio) in a growth chamber (Conviron GEN2000, 12-hour light, 80% humidity, day/night = 24/18 °C). All plants were watered weekly with deionized water; *N. klossii* clones were fertilized weekly with dilute orchid fertilizer.

### Traits of live plants

For each pitcher used in the bioassay, we measured opening diameter, lid angle, and area with fenestration. We used ImageJ (Java 1.8.0_345) to measure lid angles and opening diameters from images taken of all *N. klossii* and *C. follicularis* pitchers within 20 minutes of being cut from the plant. To maintain consistency between species, we measured the angle of opening starting at the lid apex to the point where the lid and peristome meet and then the most direct path to the outermost edge of the peristome (Figure 2.a). The opening diameter measurement consisted of the most direct path from the point closest to connection with the lid to the furthest edge of the peristome (Figure 2.b).

**Figure 2.**
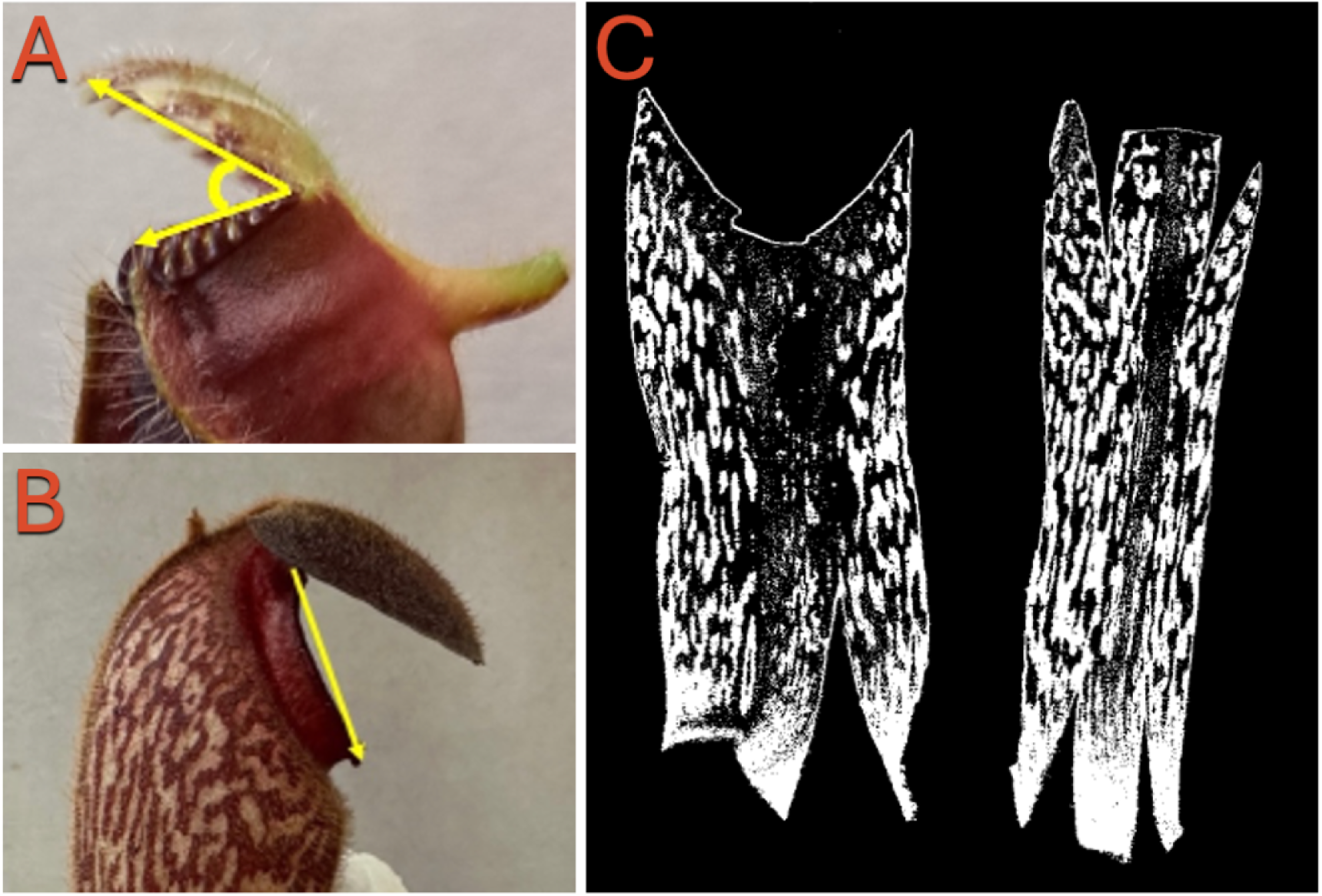
A) Schematic of ImageJ measurement of opening angle with *C. follicularis* example B) Schematic of ImageJ measurement of pitcher opening diameter with *N. klossii* example C) Binarized scan of flattened *N. klossii* pitcher with front on left side and back on right side used to quantify amount of fenestration

To measure fenestration area, we cut open, flattened, and scanned the walls and lids of all bioassay pitchers using a Canon LiDE 120 scanner. We then used ImageJ to convert the scans to binary data and quantify the percent area of fenestrated (Figure 2.c). For *N. klossii,* we compared the fenestrated area of the front and domed back of the pitcher, because fenestration occurs on the entire pitcher body, but we hypothesize a concentration of fenestration in one area is necessary for a similar light manipulation mechanic as found in *N. aristolochioides* (Moran et al. 2012). For *C. follicularis,* we compared the fenestration on the pitcher lid to the front of the pitcher, which is exposed to light, while the back of the pitcher is shaded by a dense rosette of other pitchers.

### Herbarium trait analysis

For cross-species comparison of traits that we hypothesized were related to light manipulation, we measured the lid angle and opening diameter of *N. aristolochioides* and *S. minor,* two species with previous evidence of light manipulation (McGregor et al. 2016; Moran et al. 2012). While we intended to quantify fenestration as a third trait, we were unable to consistently delineate fenestrations on scanned herbarium specimens. To compare variation in lid angle and opening diameter values between light manipulating species and close relatives, we also measured these traits from specimens of *N. singalana* which is closely related to *N. aristolochioides* (Murphy et al. 2020), *N. maxima* which is closely related to N. klossii (Murphy et al. 2020), and *S. flava,* sister to *S. minor* (Stephens et al. 2015), all species that do not manipulate light.

Images of herbarium specimens with pitcher leaves pressed in a profile were sourced from GBIF, we also used images of live pitchers from *N. singalana* and *N. aristolochioides* from iNaturalist from which we obtained only angle measurements (Wagner 2026). Six *N. aristolochioides* images taken with the same procedure as lab images were sourced from the grower Carnivero. Measurements were taken in the same way for pitchers harvested for the bioassay. To measure the angle of *S. minor*, we defined the vertex of angle as the topmost part of its fenestration, since it has a hooded morphology.

### Scanning Electron Microscopy (SEM)

To examine the pitcher’s microscopic structures for convergence across our study species, we imaged fresh specimens using a Hitachi TM3030 microscope to examine the inside back wall of the pitcher for antiadhesive wax crystals and the peristome for a wettable, anisotropic surface (Bohn and Federle 2004; Thorogood et al. 2018).

### Fly bioassay

To assess whether light entering fenestrated areas influences overall prey capture, bioassays were conducted in an isolated system exposing live flies to freshly cut pitchers with and without light blocked (Moran et al. 2012). The bioassay apparatus consisted of a 1-liter clear plastic container, a paper stand for the cut pitcher to sit on, red plastic filter paper attached to a wire base, and a full spectrum light above (Reptisun 10.0 UVB). For prey, we used lab-reared *Drosophila suzukii* flies as a model for small flying Diptera species captured by *N. klossii* in its native range (Moran et al. 2012). *Drosophila* flies have limited sensitivity to longer wavelengths of light such as red and infrared (Little et al. 2019; Sharkey et al. 2020), so we used a red plastic filter (650 nm reflectance peak, Online Resource 1) to block visible wavelengths from entering fenestrations, negating the pitcher’s ability to manipulate light and disorient the flies in treatment trials. To isolate the effect of light manipulation from any effect of the red filter itself, we included the red filter in control trials but distanced it from the pitcher and angled it away from fenestrated areas. Individual pitchers were harvested immediately before bioassays began and were not reused. Before the trials began, all flies were starved for approximately 12 hours with access to a moist paper towel. Ten flies were added to each container containing the pitcher and filter, and after three hours we scored the survival of each fly to calculate the total capture (*N. klossii* N = 16, *C. follicularis* N = 20).

### Statistical analysis

To analyze our data, we used R Studio version 2025.09.2+418 and the car, performance, glmmTMB, and DHARMa packages (Bolker 2020; Fox et al. 2013; Hartig 2020; Lüdecke et al. 2021). To compare the difference in proportion of flies captured between the control and treatment trials within each species, we used Beta Generalized Linear Models (GLMs) with treatment as a fixed effect. We did not include plant as random effect because, within each species, all plants were genetic clones grown under identical conditions in a growth chamber and thus were treated as a single pool. To examine the relationship between our proposed component traits and fly capture, we used a Beta GLM for both species with lid angle, pitcher opening, and fenestration difference as fixed effects. To compare the fenestration on the front and back of *N. klossii,* we used a Beta GLMM to model the percent area fenestrated, with location as a fixed effect and leaf ID as a random effect. We used a Levene’s test to compare the trait variation between each pair of closely related species for opening diameter and lid angle.

## Results

### Fenestration Measurements from Experimental Plants

Quantification of *N. klossii* and *C. follicularis* fenestrations demonstrated both species have significant concentrations of fenestrations on dorsal surfaces where light could pass through into the pitcher interior. In *N. klossii,* evidence showed the dorsal side of the pitcher had 122% [95% CI: 104, 142] more fenestration than the ventral side (Figure 3). The back of *N. klossii* pitchers had an average of 46% percent apigmented area compared to only 28% apigmented area on the front (p < 0.0001). We found that *C. follicularis* lids on average were 26.6% fenestrated, compared to no fenestration on the front wall of the pitcher (Figure 3).

**Figure 3.**
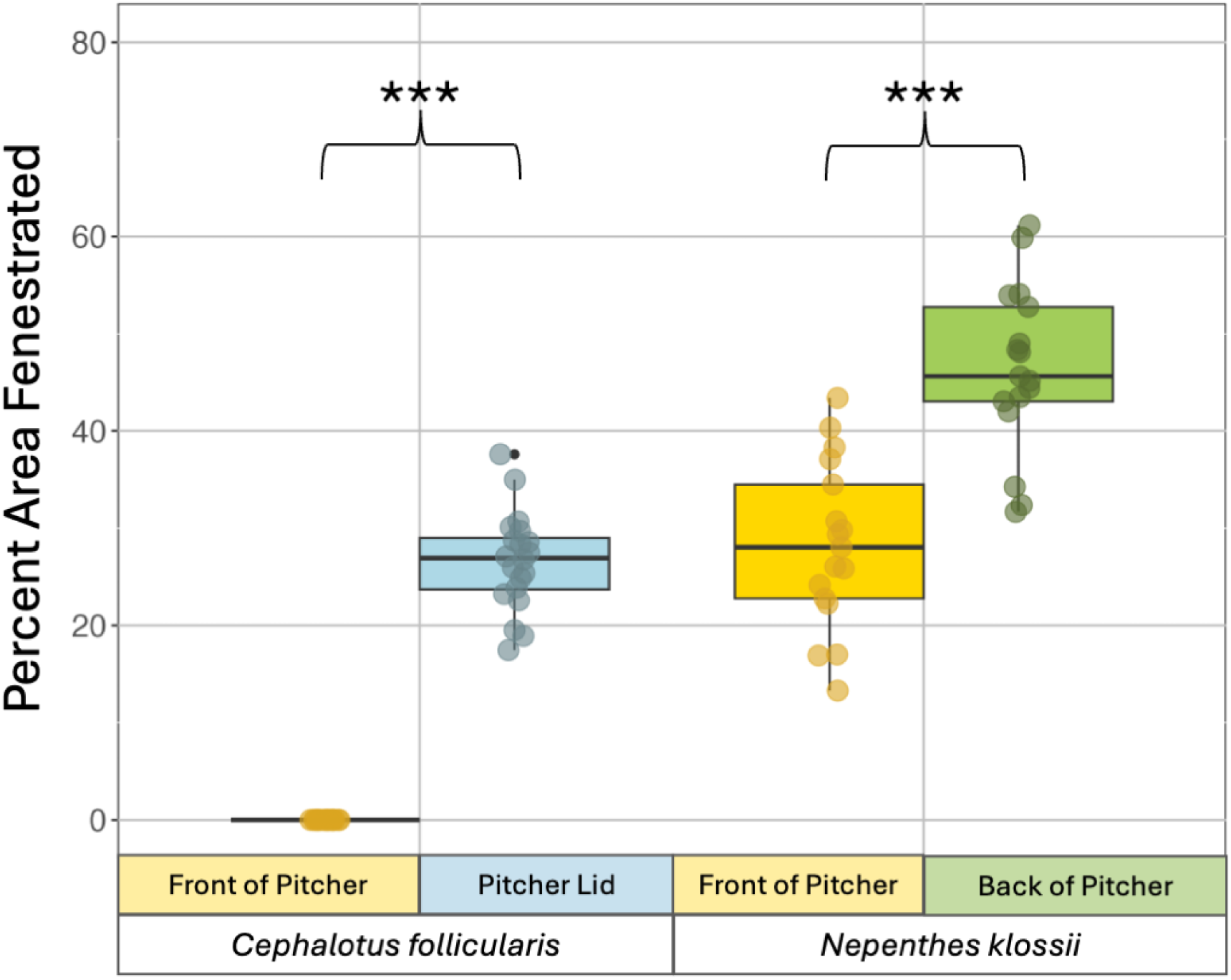
Both *C. follicularis* (N = 24) and *N. klossii* (N = 20) show a significant amount of fenestration in distinct regions where light passes through the pitcher, meaning incoming light is concentrated and may be a point of false exit for prey.

### Trait Variance Results

Proposed component trait values of *N. klossii* and *C. follicularis* were strikingly similar to *N. aristolochioides* and *S. minor* (Figure 5). Using a Levene test, we found that overall, the selected species with fenestrations have significantly narrower variances in lid angle and opening diameter compared to closely related species, as well as similar ranges to *N. aristolochioides.* When comparing *S. minor* to *S. flava, S. minor* had a significantly smaller variance in opening diameter (P < 0.004) and lid angle (P < 0.008). *N. klossii* showed significantly smaller variances in opening diameter (P < 0.005) and lid angle (P = 0.003) than *N. maxima*. Between *N. aristolochioides* and *N. singalana, N. aristolochioides* had a significantly smaller opening diameter (P = 0.02), but not a significantly smaller lid angle (P = 0.76). Since *C. follicularis* is the only pitcher species in its order, there is no related species as a direct point of comparison for its trait variances, however it shows similar trait values to both *N. aristolochioides* and *S. minor* (Figure 5). When comparing only among trait measurements from herbarium records, we see the same trends of lower variance in lid angle and opening diameter for fenestrated species, so we are confident that this result is not driven by trait measurement methodology between live *N. klossii* and *C. follicularis* images, pictures from carnivorous plant growers, iNaturalist observations, and herbarium records.

### SEM Results

Unlike its relative *N. aristolochioides,* SEM images of the inner back wall of *N. klossii* show it retains antiadhesive waxy crystals (Online Resource 5). *C. follicularis* does not possess wax crystals on its inner pitcher walls (Adams and Smith 1977). The peristome of the *N. klossii* and *C. follicularis* both have an anisotropic surface structure (Online Resource 5) that can cause the surface to be extremely slippery when wet (Figure S6, Bohn and Federle 2004).

### Fly Bioassay and Component Traits

In contrast to previous experiments using *N. aristolochioides* (Moran et al. 2012), the red filter blocking visible light did not have a significant effect on the number of flies captured by *N. klossii* or *C. follicularis*. *N. klossii* pitchers with a red filter covering their fenestrations caught 68% of flies compared to pitchers without the filter that caught 73% although this difference was not significant (P = 0.5498, Figure 4). *C. follicularis* pitchers that had a red filter covering their fenestrations caught 65% of flies while *C. follicularis* pitchers without a red filter caught 64%. (P = 0.857, Figure 4).

**Figure 4.**
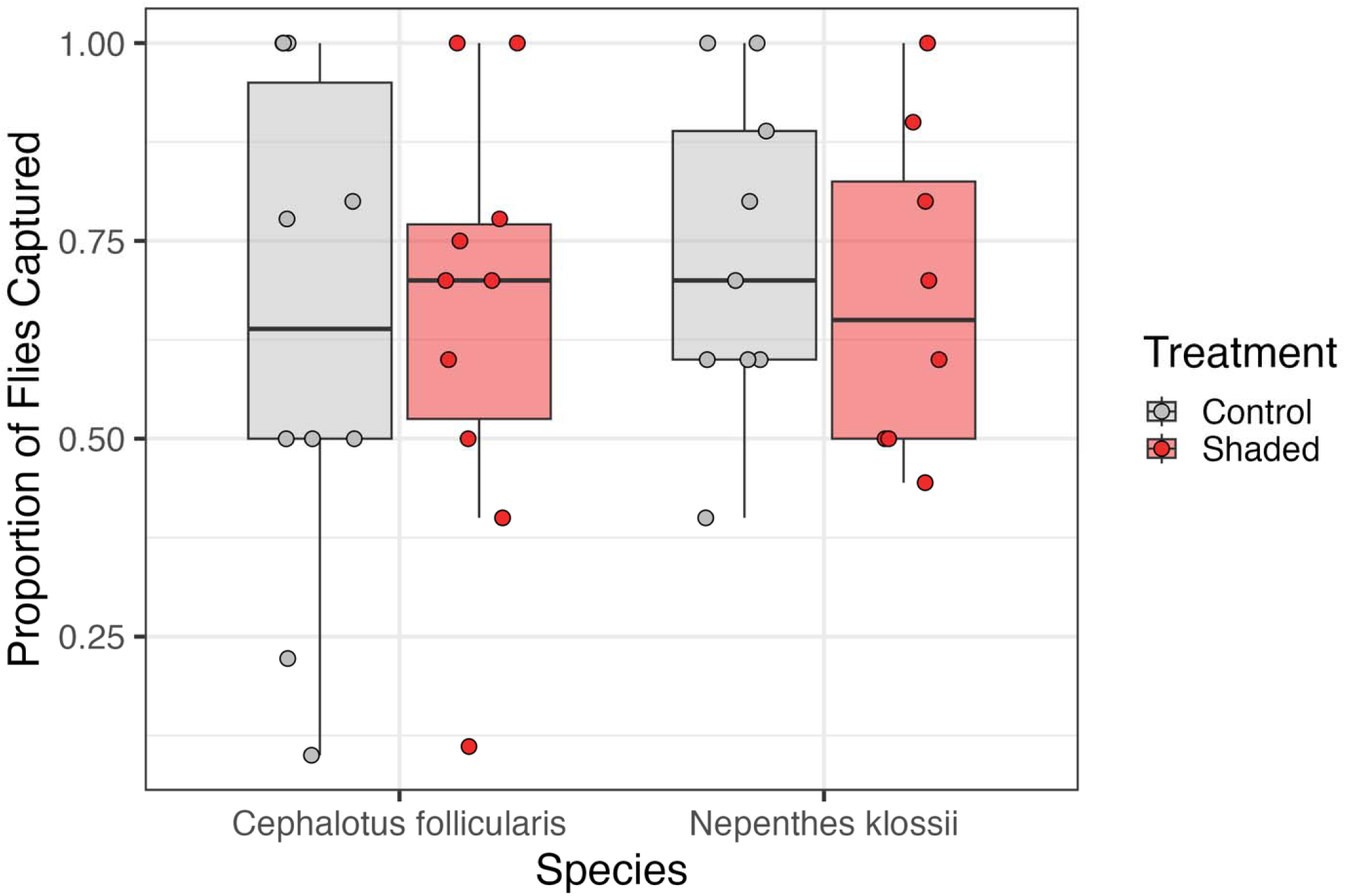
The shaded treatment had a nonsignificant positive effect on fly capture for *Cephalotus follicularis* (total N in experiment = 19) and a nonsignificant negative effect on fly capture for *Nepenthes klossii* (total N = 17).

**Figure 5.**
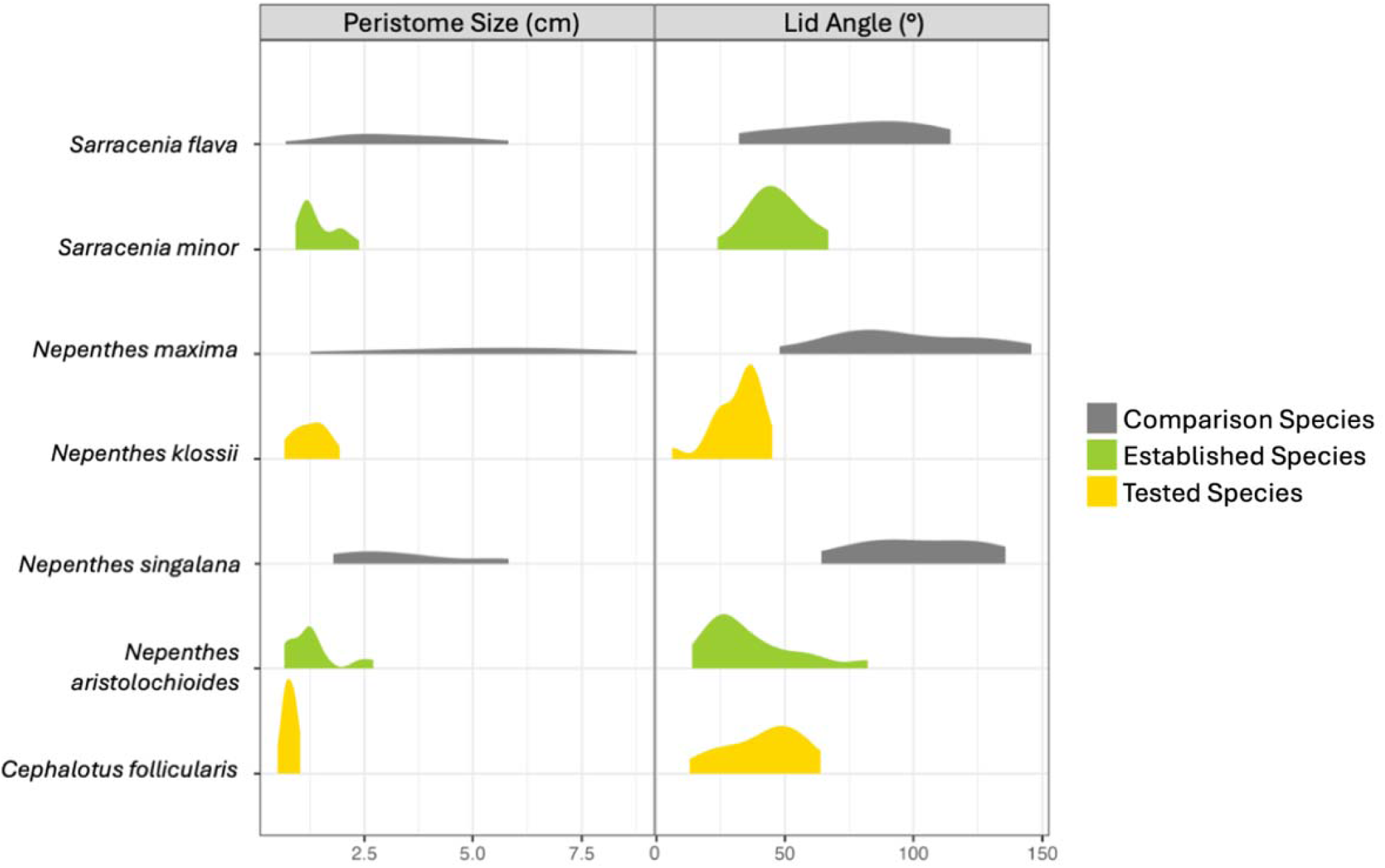
In species proposed or demonstrated to manipulate light, component trait ranges and variances are more similar to each other than to closely related species that do not manipulate light.

Pitcher opening diameter for *N. klossii* was the only component trait that significantly influenced the amount of prey captured for either species. For every 1 mm increase in opening diameter width for *N. klossii*, there was a 14.2% decrease in the odds of prey capture (95% CI [3.02, 24.13], P = 0.008, Online Resource 2), consistent with our hypothesis that smaller pitcher openings may increase the effectiveness of light manipulation. However, this relationship was nonsignificant and positive for *C. follicularis* (P = 0.351, Online Resource 2). A larger lid angle had a negative, nonsignificant relationship with fly capture for *N. klossii,* although the nonsignificant relationship was positive for *C. follicularis*, in opposition to our expectation for a light manipulating species (Online Resource 3). There was a nonsignificant negative relationship between an increase in the difference in fenestration and fly capture for both species (Online Resource 4).

## Discussion

In this work, we tested if two convergently evolved pitcher plant species (*Nepenthes klossii* and *Cephalotus follicularis)* increase prey capture by manipulating light and quantified the variances of component traits by integrating bioassays, live plant and herbarium measurements, and scanning electron microscopy. Both species showed similar patterns of fenestration to light-manipulating species, as well as similar proposed component trait values (opening diameter and lid angle) to *N. aristolochioides,* a light-manipulating species. However, our lab bioassay did not support our hypothesis that *N. klossii* and *C. follicularis* increase prey capture by manipulating light through their fenestrations, though a smaller opening diameter did predict an increase in prey capture for *N. klossii* (Online Resource 2). While additional laboratory and especially field experiments may reveal a light manipulation function too weak to be detected in our study, together, our results may point to convergent morphology in these two species arising from selection acting on a different function beyond light manipulation for prey capture.

Fenestration may provide an increase to pitcher plant fitness for a reason unrelated to prey capture. One hypothesis for the function of fenestration is that it plays a role in long-range prey attraction. Some work has found that covering fenestrations decreased the visitation of insects rather than capture of prey for *S. minor* (Schaefer and Ruxton 2014). This may be the case particularly for *C. follicularis,* since the location of its fenestrations is conspicuous from above instead of opposite from the pitcher opening like other fenestrated pitcher species, and may provide visual contrast to attract flying insects rather than confuse them with false exits.

Alternatively, work in noncarnivorous plants has found that bundle sheath extensions in fenestrations allow more light into internal layers of dense leaf which increases photosynthetic performance (Karabourniotis et al. 2021). Though many species in the *Nepenthes* family have low photosynthetic performance in their pitchers compared to their lamina, or leaf blades, *C. follicularis* pitchers have similar proportions of light absorption to their lamina, and surprisingly their pitchers have similar photosynthetic rates to pitcher species like *Darlingtonia californica* that both trap and photosynthesize with their pitchers (Pavlovič 2011). Further experimentation with *C. follicularis* and *N. klossii* would be beneficial in exploring these hypotheses.

A key difference between the prey capture strategies of *N. aristolochioides* and *N. klossii* explaining the observed difference in light manipulation function may be their pitchers’ fluid and microstructure. The walls of *N. aristolochioides* are coated in a highly viscoelastic fluid that is nearly impossible for prey to escape from (Gaume and Forterre 2007). Moran et al. (2012) describe a hypothesized capture scenario in *N. aristolochioides* in which flies enter the pitcher and attempt to leave via a false exit (fenestration) and become ensnared upon contact with the pitcher wall where viscoelastic fluid is present. In contrast, flies may readily escape after dropping into more aqueous *Nepenthes* fluid that is only found at the bottom of the pitcher (Gaume and Forterre 2007). *N. klossii* fluid much less viscoelastic than *N. aristolochioides* which may make it less efficient at ensnaring flying prey, though *N. klossii* is still stickier than water-like *C. follicularis* fluid (ESW, pers. obs.). Instead of *N. aristolochioides’* sticky walls, we found that *N. klossii* has wax-covered walls, which may attenuate light coming through fenestrations to a greater degree than only a fenestrated area (Moran et al. 2012), though a difference in wax between upper and lower pitchers is possible (Gorb 2009). There is evidence to support a macroevolutionary tradeoff between wax and viscoelasticity, with viscoelastic fluid shown to be most effective for fly capture (Bonhomme et al. 2011). *N. klossii* and *C. follicularis* do not possess strongly viscoelastic fluid, and this may explain why we observed a fenestration difference, low lid angle, and narrow opening diameter similar to *N. aristolochioides* and *S. minor* but not a detectable light manipulation function for either *N. klossii* or *C. follicularis* in the fly bioassay. Future studies could manipulate the presence of viscoelastic fluid and wax on the pitcher wall to determine the criticality of this trait to the light manipulation strategy across species.

Additional pitcher morphological differences may contribute to the lack of light manipulation demonstrated in *C. follicularis* and *N. klossii*. Older upper canopy *N. klossii* pitchers have a pronounced domed morphology that more closely resembles *N. aristolochioides* morphology (McPherson 2009), but the pitchers used in our bioassays were younger lower pitchers that were less domed. Like a convex lens (Hecht 2017), a more dramatically domed pitcher morphology may direct incoming sunlight toward a single point inside the pitcher after coming in contact with the window-like fenestrations, concentrating the light and enhancing the false exit effect. Extreme domed morphology is present in both *N. aristolochioides* and *S. minor,* as well as other species hypothesized to manipulate light. In addition, SEM results showed that *N. aristolochioides* and *N. klossii* both have anisotropic epidermal cells on their peristomes that make the pitcher rim extremely slippery when wet (Bonhomme et al. 2011). This is a widespread trait across *Nepenthes,* but one that may not be beneficial in *N. klossii* because it has a nearly vertical peristome and may also specialize on flying prey, like *N. aristolochioides*; however, the main prey items of *N. klossii* are poorly characterized, and a wettable peristome could be adaptive if crawling prey could comprise a significant part of the species’ diet.

The similarity in the range and variance of proposed component trait values between *N. aristolochioides*, which is known to manipulate light (Moran et al. 2012), and *N. klossii* and *C. follicularis*, is striking. There are three explanations for reduced variance in opening diameter and lid angle in these species in addition to the null statistical expectation that trait distributions with a lower mean will also have a lower variance: 1) selection is unrelated to light manipulation, and is either pleiotropic or related to a different function. Indeed, with a small bioassay sample size, only a smaller opening diameter for *N. klossii* significantly increased prey capture out of all the component traits. Alternatively, 2) drift can reduce variation in the absence of selection. While this is a possibility, we find selection a more likely explanation than drift for the reduced trait variance we observed. While some species have small range sizes and thus likely small population sizes, range size does not correlate with possible light manipulation function. Finally, 3) a light manipulation function could be expressed very weakly and is undetectable in our study. If so, it would imply light manipulation is much weaker in these species than in *N. aristolochioides*, which indeed does have more extreme domed morphology and a greater difference in concentration of fenestration than either *N. klossii* or *C. follicularis* (Moran et al. 2012). A small marginal increase in capture rate could exert substantial selection on component traits over evolutionary time, yielding low component trait variances but undetectable light manipulation function.

Our data are consistent with a directional selection scenario of complex function evolution. In a directional selection scenario, component trait phenotypes shift towards the trait values required for complex function, due to selection from weak functioning of the complex trait, selection on the trait for a separate function, or pleiotropic selection. In a spontaneous coincidence scenario, the specific values of component traits required for complex function spontaneously co-occur and cause a significant increase in fitness to the organism. This implies independent evolution of each trait and that species without the complex function would have greater phenotypic variation in comparison to species that do. In both *N. klossii* and *C. follicularis*, it appears that the traits of fenestration, opening diameter, and lid angle have undergone selection without a light manipulation function present, consistent with selection acting on these traits before a complex function arises. This differs from the findings of Chomicki et al. (2024) on the evolution of the complex function of springboard trapping in two distantly related *Nepenthes* species, which found evidence consistent with a spontaneous coincidence rather than directional selection scenario of complex function evolution.

## Conclusion

In this study, we asked if fenestrations in pitcher plants worked in combination with other component traits to allow for a unique prey capture mechanism as found in *N. aristolochioides* (Moran et al. 2012). While many component traits had similar ranges and variance to those of *N. aristolochioides*, light manipulation was not detected in in both *C. follicularis* and *N. klossii*.

While this study does not offer a definitive explanation for the convergence of fenestrations in pitcher plants, it does highlight the complexity of untangling morphological function reliant on a suite of traits and the specific functionality of component traits across different lineages.

## Acknowledgements

We thank Dr. Monica Mowery, Dr. Marianna Szucs, Dr. Julianna Wilson for consult on and supply of fly cultures; the Bronikowski lab and KBS facilities for necessary tools; Rosemary Glos for SEM imaging assistance; and Carnivero for pitcher plant measurements and supply of plants.

## Declarations

### Ethical approval

All applicable institutional and/or national guidelines for the care and use of animals was followed.

### Consent to participate

Not applicable.

### Consent for publication

Not applicable.

### Availability of data and material

The data was deposited on Dryad under the reference number 10.5061/dryad.cvdncjtkm which will become live upon manuscript acceptance. The link for private peer review is: http://datadryad.org/share/LINK_NOT_FOR_PUBLICATION/9O18XyqjiaMC_F5dpNuhTjoagnm2tWfWOv2aRKfq9u4

### Code availability

The code was deposited on Dryad under the reference number 10.5061/dryad.cvdncjtkm which will become live upon manuscript acceptance.. The link for private peer review is: http://datadryad.org/share/LINK_NOT_FOR_PUBLICATION/9O18XyqjiaMC_F5dpNuhTjoagnm2tWfWOv2aRKfq9u4

### Highlighted Student Paper Consideration

This study should be a Highlighted Student Paper because of its significance as a multi-faceted approach to investigating selective processes on complex function in organisms. Plant behavior is currently a major frontier of knowledge in ecology, and this work makes cross genera comparisons linking convergent morphologies to potential fitness increase in charismatic carnivorous plants. This paper demonstrates a successful framework to quantify component traits that has wide application in ecology and evolution.

### Author’s contributions

ESW and SME conceived and designed the experiments, performed experiments, collected data, analyzed the data, and wrote the manuscript; other authors provided editorial advice.

