## Supplement for "Shedding light on fenestrations in carnivorous pitcher plants: a test for convergent function"

ELECTRONIC SUPPLEMENTAL MATERIAL

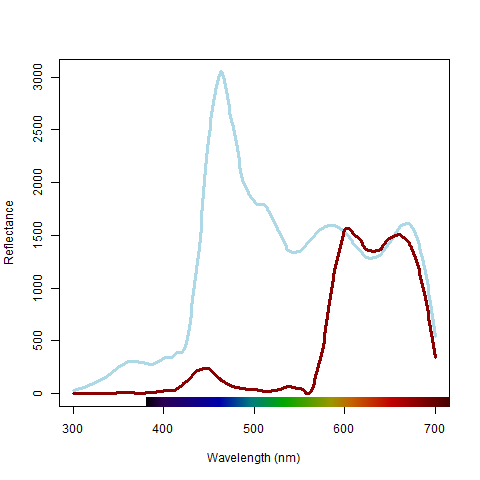

**Online Resource 1.** Blue line shows full spectrum reflectance from Reptisun 10.0 UVB. Red line shows reflectance from red plastic used as filter for bioassays, which had lowered reflectance in wavelengths of light visible to *Drosophila suzukii.*

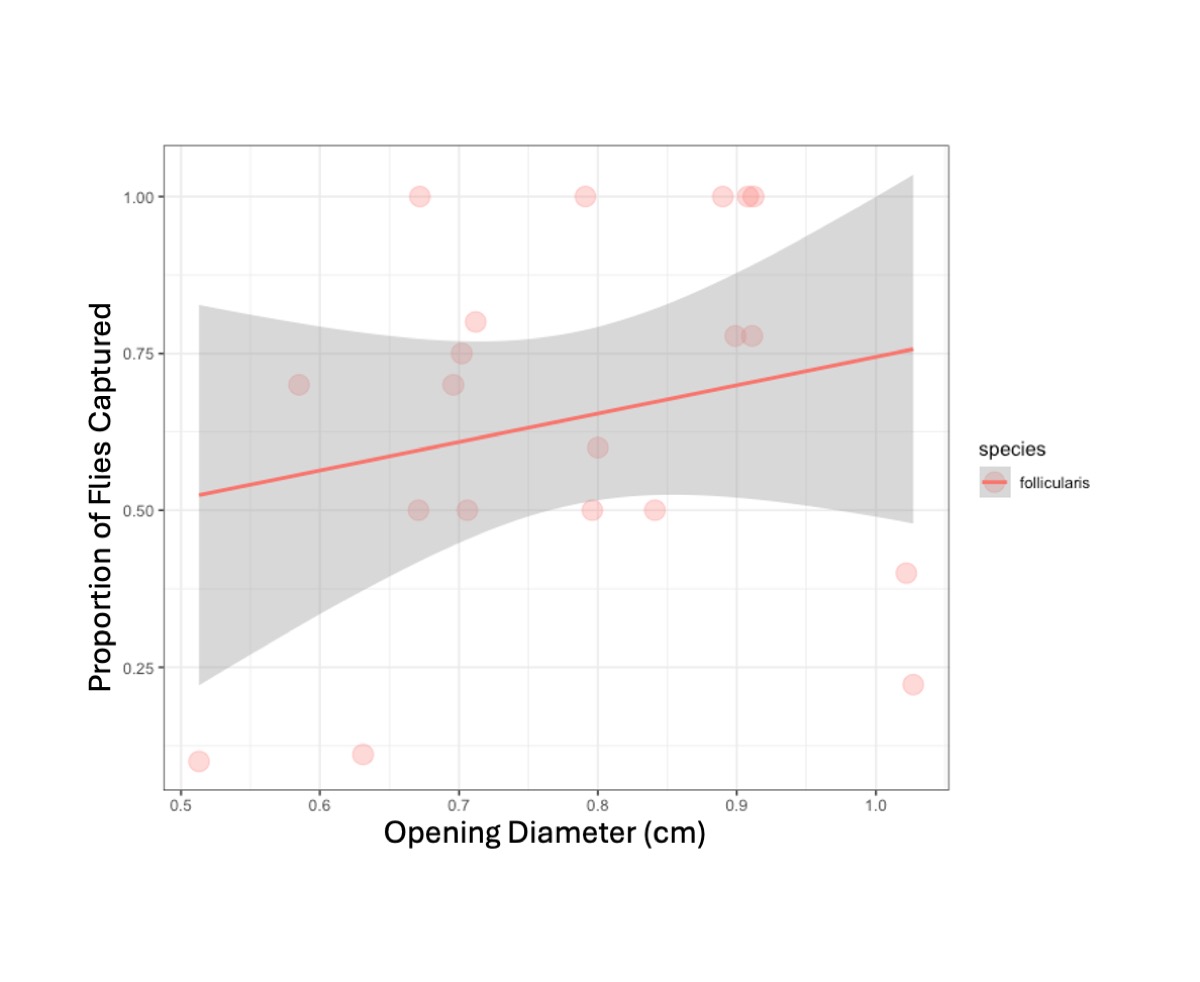

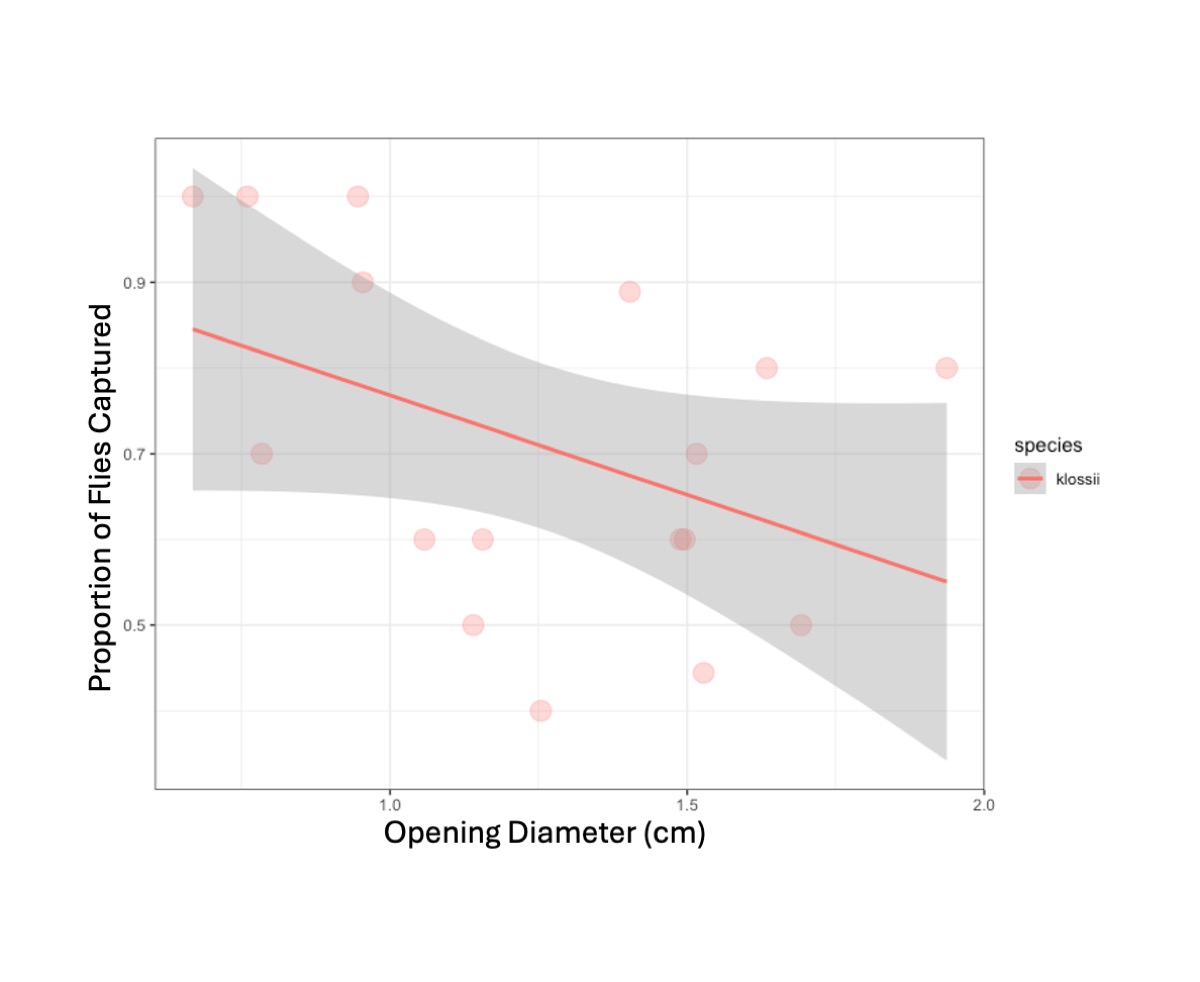

**Online Resource 2.** The increase in the size of the pitcher mouth opening had a positive, insignificant effect on fly capture for *C. follicularis* (P = 0.351) and a negative, significant effect on fly capture for *N. klossii* (P = 0.008).

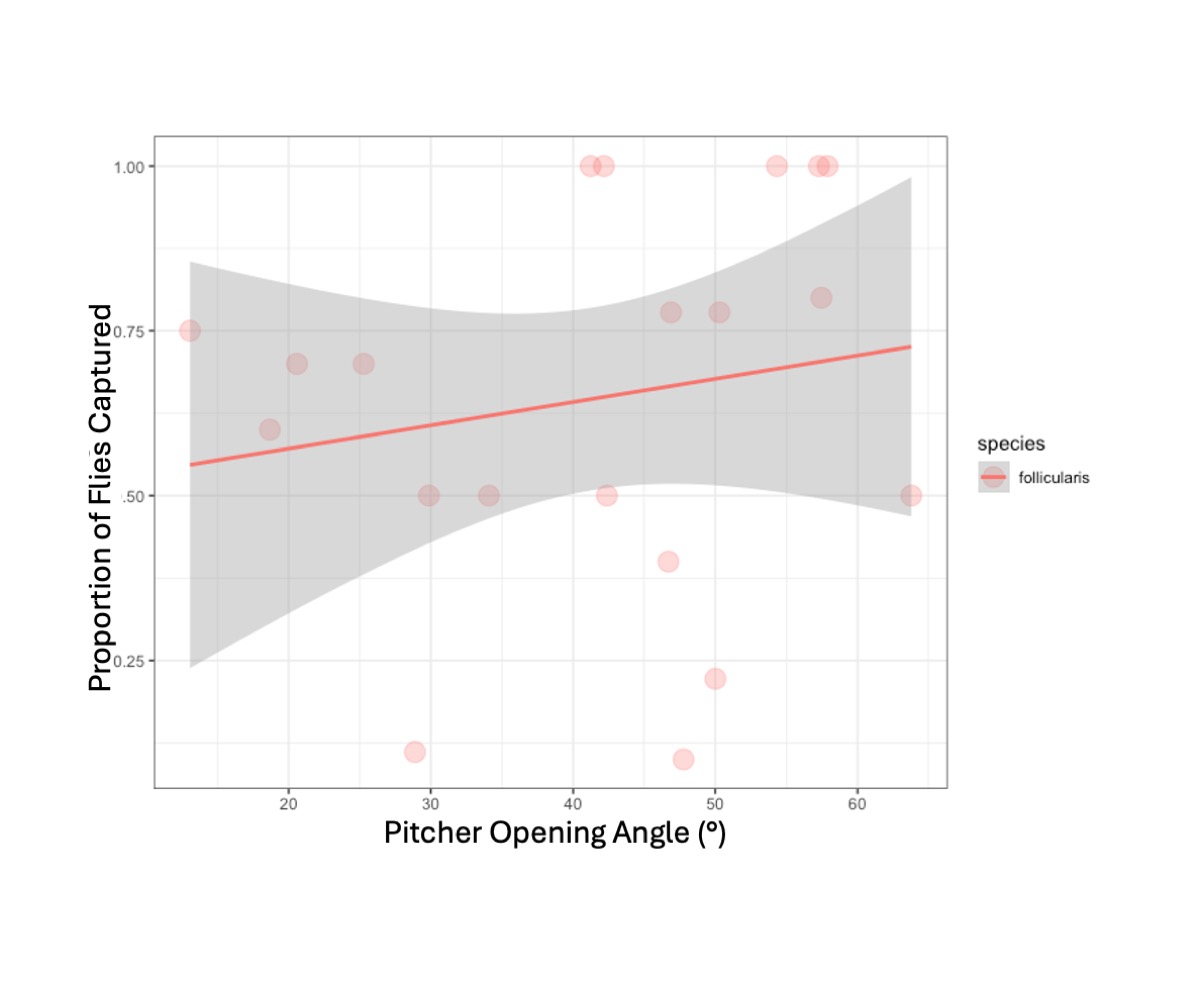

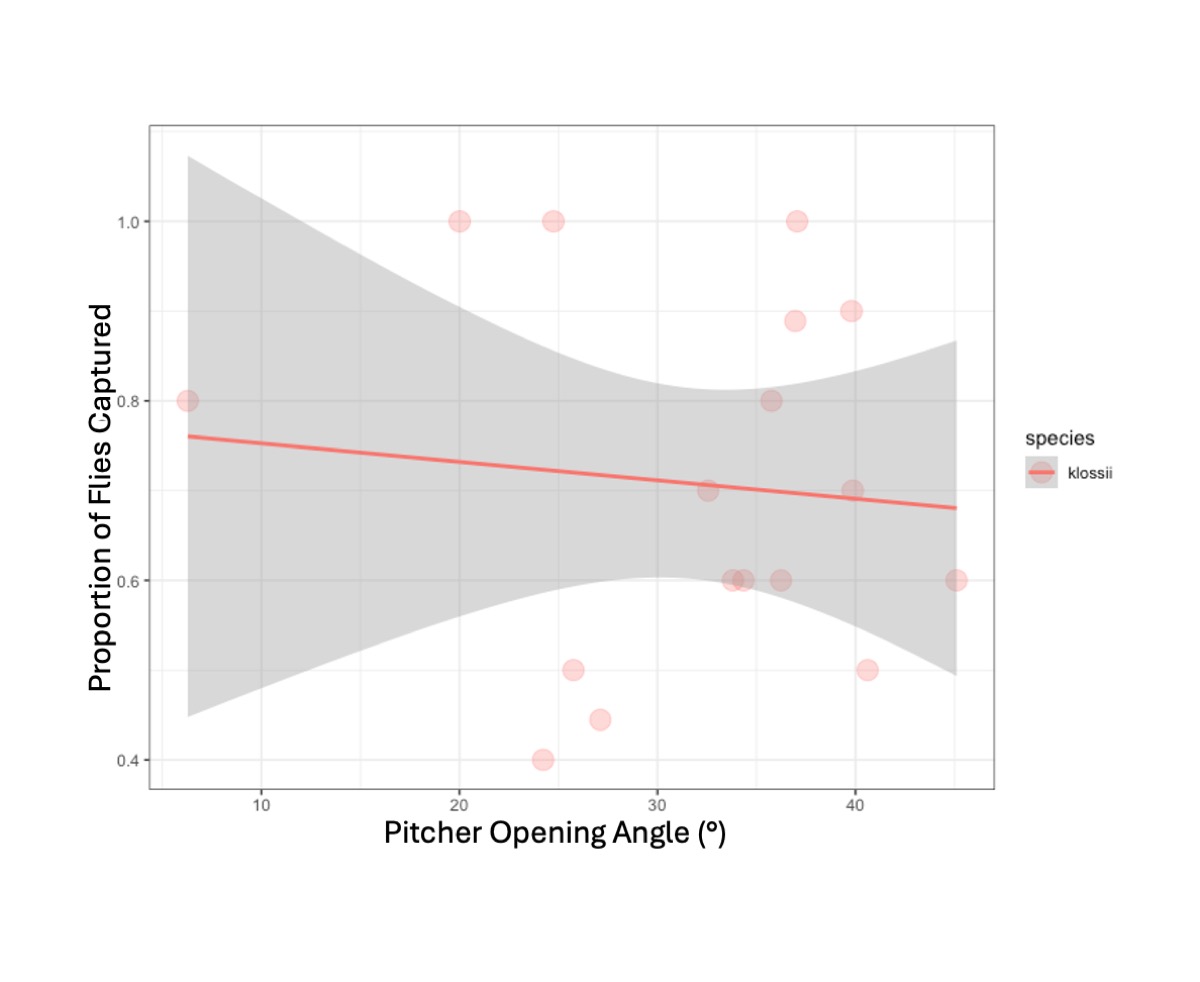

**Online Resource 3.** The increase in the angle of the lid to the pitcher mouth had a positive, non-significant effect on fly capture for *C. follicularis* and a negative, non-significant effect on fly capture for *N. klossii.*

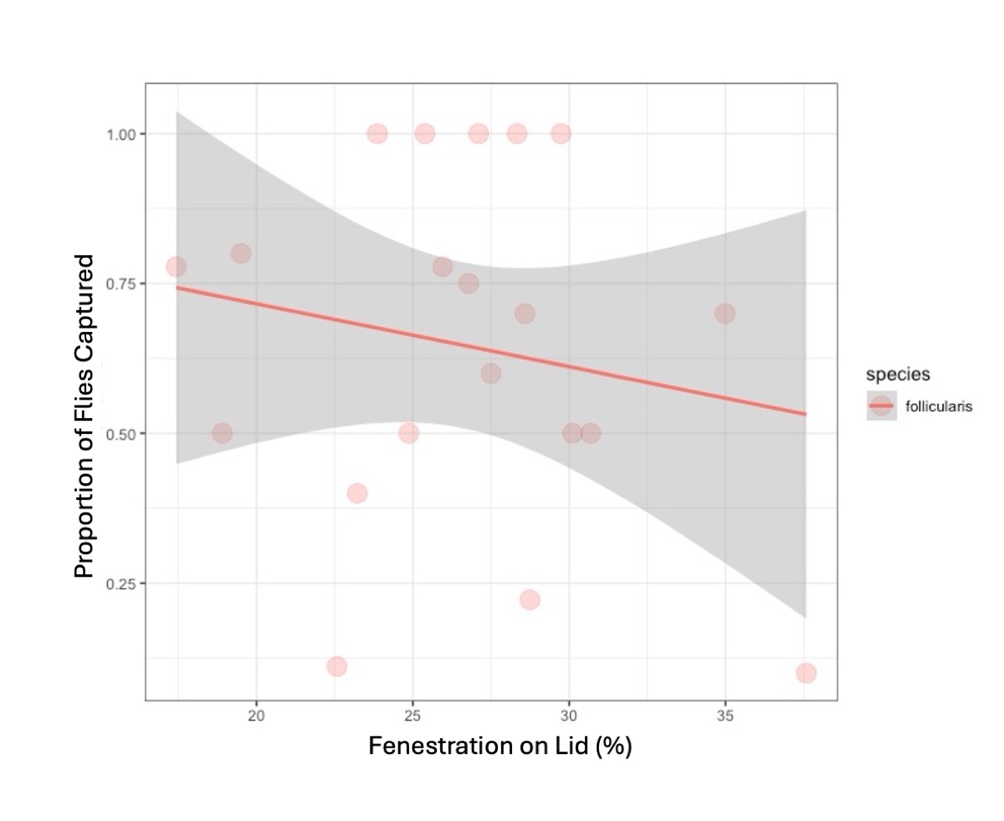

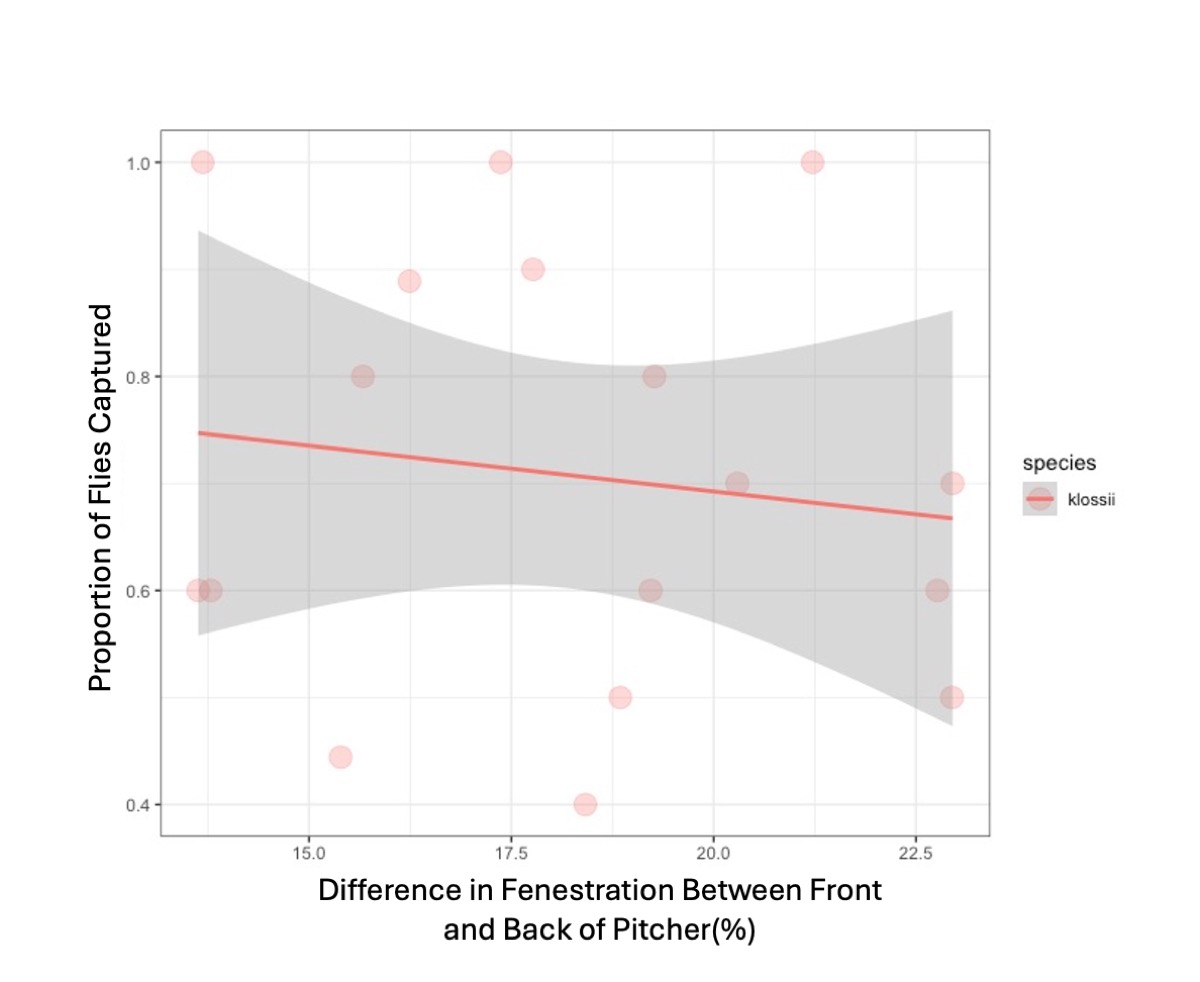

**Online Resource 4.** The increase in the percentage of fenestrated area had a negative, non-significant effect on fly capture for *C. follicularis* and *N. klossii.*

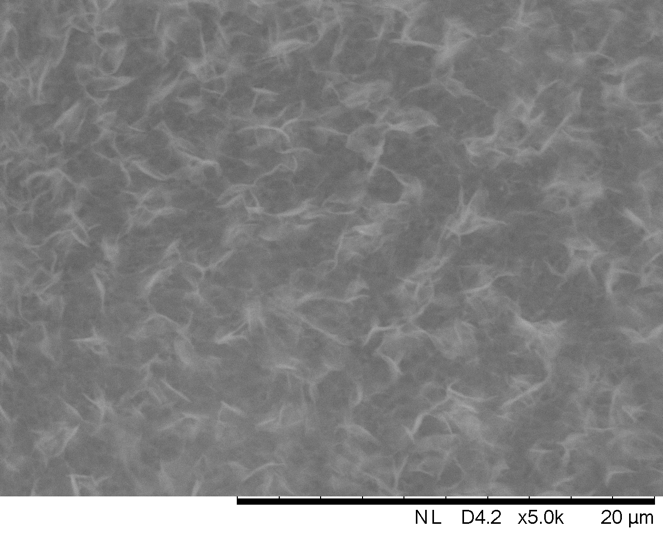

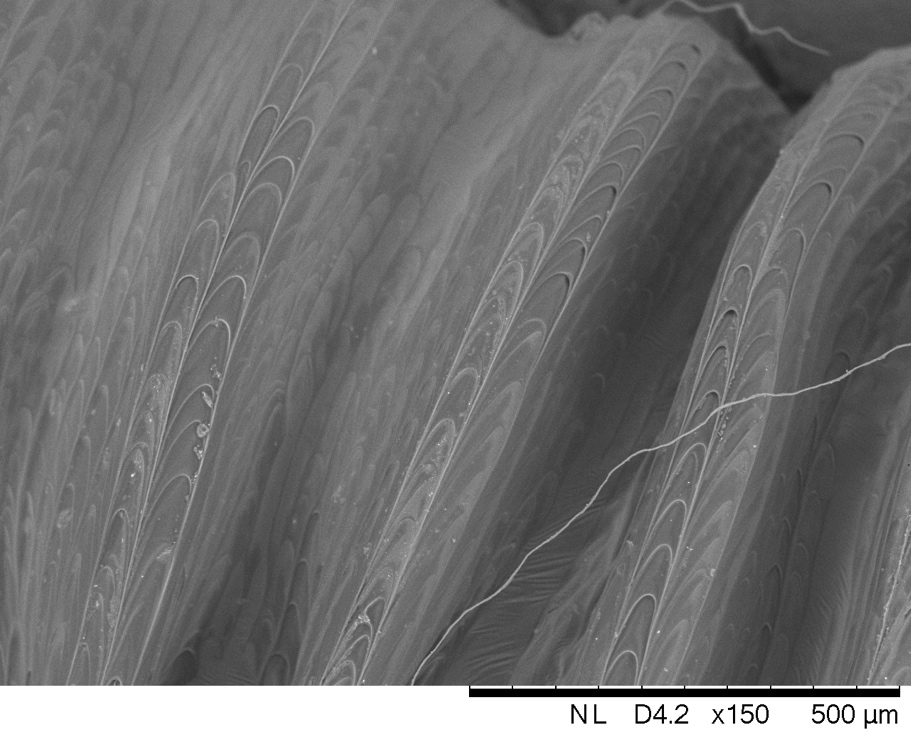

**Online Resource 5.** Scanning electron microscopy results shows (Top) antiadhesive wax crystals on the inside of *Nepenthes klossii* pitcher and (Bottom) anisotropic structure on *N. klossii* peristome, previously shown to make surface wettable and slippery when wet for insects.
